# Estimating HLA disease associations using similarity trees

**DOI:** 10.1101/408302

**Authors:** Christiaan H. van Dorp, Can Keşmir

## Abstract

The human leukocyte antigen (HLA) is associated with many (infectious) disease outcomes. These associations are perhaps best documented for HIV-1. For example, the HLA-B*58:01 allele is associated with control of the virus, while HLA-B*18:01 is considered detrimental. In HLA disease association studies, it is often ignored that certain HLA molecules are functionally very similar to others. For instance, HLA-B*18:03 differs “only” at 3 positions in its peptide binding site from HLA-B*18:01, and not surprisingly, HLA-B*18:03 is also associated with fast progression to AIDS. Here, we present a Bayesian method that takes functional HLA similarities into account to find HLA associations with quantitative traits such as HIV-1 viral load. The method is based on the so-called phylogenetic mixed model (a model for the evolution of a quantitative trait on the branches of a phylogeny), and can easily be modified to study a wide range of research questions, like the role of the heterozygote advantage, or KIR ligands on disease outcomes. We show that in the case of HIV-1, our model is significantly better at predicting set-point virus load than a model that ignores HLA similarities altogether. Furthermore, our method provides a comprehensible visualization of HLA associations. The software is available online at www.github.com/chvandorp/MHCshrubs

## 1 Introduction

The major histocompatibility complex (MHC) molecules, are key elements of the cellular adaptive immune system. MHC molecules present peptides to T cells, triggering an immune response when the T-cell receptor (TCR) binds the peptide-MHC (pMHC) complex. Pathogens can escape from T-cell responses by mutating amino-acids in their peptidic epitopes, thereby disrupting the MHC-peptide binding, or the pMHC-TCR binding. It has been suggested that in order to cope with the first kind of escape, massive MHC polymorphism exists on the population level [41, 25]. This polymorphism ensures that pathogens are unable to escape from every immune response, since functionally, MHC polymorphism results in differential peptide presentation. Moreover, many MHC molecules are ligands for Killer-cell immunoglobilin-like receptores (KIRs) on natural killer (NK) cells, increasing the the pathogen-driven selection on MHC molecules even further.

The human MHC molecules, human leukocyte antigens (HLA) have been associated with many different disease outcomes, both for auto-immune diseases as for infectious diseases. Some notable examples are the HLA class II alleles HLA-DQ2 and HLA-DQ8 which are strongly associated with coeliac disease [40], and the class I allele HLA-B27, which has been associated with a variety of auto-immune diseases, such as ankylosing spondylitis [46]. Interestingly, the causal link between HLA-DQ2 and DQ8 and coeliac disease is well understood [40], while the mechanism that causes the HLA-B27 disease associations is not known, although several theories have been formulated [46].

Perhaps the best studied example of associations between HLA alleles and infectious disease traits is HIV-1 infection [27]. For instance, HLA-B57 and HLA-B27 are enriched in so-called long-term non-progressors: patients that have asymptomatic infections for a very long time and a low set-point virus load (SPVL). Other alleles have been associated with a high SPVL and are considered detrimental [15, 19]. HLA effects on SPVL are relatively well understood [27], especially in the case of protective alleles, where many immune responses against specific HLA-restricted epitopes have been identified [4]. Especially responses against epitopes in the Gag polyprotein are associated with viral control. Furthermore, both the within-host and population-level evolution of HIV-1 has been shown to be driven by HLA [17], which can lead to the loss of HLA-SPVL associations over time [37, 3].

Finding associations between HLA and disease traits is difficult for a number of reasons. First and foremost, the number of HLA molecules is very large. This results in statistical models with many parameters, and a serious risk of over-fitting. To prevent this, some kind of regularization (i.e., imposing Occam’s razor) must be invoked [5], or a stringent multiple-testing correction is applied [30, 15]. Secondly, but related to the first issue, the allele frequency distribution is far from uniform. In European populations, the most common allele of the A locus (HLA-A*02:01) has a prevalence of 30%, but most alleles are very rare [34]. Therefore, large sample sizes are required in order to gather enough observations involving these rare alleles. In some studies, alleles that are e.g. sampled less than 5 times are left out of the analysis, since any effect found for these alleles is not meaningful statistically. Moreover, leaving out alleles reduces the complexity.

Thirdly, different HLA molecules are evolutionary and functionally related, and should therefore not be considered as independent entities in an association study [13, 41]. The relatedness between HLA molecules is partially captured by the nomenclature (e.g. A*02:01), where the first field (02) is used to designate the allele type, which is based on genetic similarity, and the second field (01) determines the subtype. In some association studies, alleles of the same type are lumped together. This partially solves the issues discussed above as: (i) The number of HLA types is much smaller than the number of subtypes. (ii) Rare alleles are grouped together with more common alleles of the same type, which increases the number of observations per covariate, and avoids wasting observations. (iii) Molecules of the same type presumably are functionally more similar than molecules of two separate types, and hence, HLA type can be regarded more accurately as independent entities than HLA subtypes.

Unfortunately, using HLA types for association studies instead of subtypes has some major limitations. Most importantly, the effect of HLA on disease can be on the subtype level and not on the type level. For example, in the case of HIV-1, the allele HLA-B*58:01 is known to be protective, while its ‘cousin’ HLA-B*58:02 is detrimental. Moreover, some types are closely related to other types (e.g. HLA-B*57 and HLA-B*58), making the use of types both too course grained and not course grained enough for solving issue (iii) above. Some authors therefore choose to aggregate alleles in some types (such as B*57), but treat other alleles individually (e.g. B*58:01 and B*58:02), in a rather ad hoc fashion [15, 19]. Another method that has been used previously [5], makes use of HLA supertypes [38, 21] for the aggregation of HLA alleles, which are groups of HLA alleles based on peptide binding properties. Grouping HLAs within supertypes has similar shortcomings as the approach using HLA types [43].

In this paper, we present a new method that addresses the above-mentioned issues in a more sophisticated manner. Our method uses prior information about the functional relatedness between HLA molecules based on the amino acid sequence. Therefore, we can estimate associations between ‘clades’ of HLA alleles, as well as individual alleles and a disease trait simultaneously. Because the method allows for the inclusion of prior information, it is natural to formulate this statistical model in a Bayesian framework. The benefits of our methods include those that come from grouping alleles: rare alleles can be included in the analysis, and the Bayesian framework provides natural regularization of the parameters.

We test our method on a publicly available data set of HIV-1 SPVL and HLA class I genotypes [15]. In order to construct a HLA-relatedness tree, we primarily use so-called HLA pseudo-sequences consisting of 34 non-consecutive amino-acids in the peptide binding groove of the HLA protein, that are variable across different molecules [10]. Statistically, our method is better at predicting SPVL from the HLA genotype than a model that assumes independence between HLA alleles. Finally, we demonstrate the flexibility of our model by exploring several biological scenarios, such as the role of HLA as a ligand for natural killer (NK) cells, and the effect of homozygous HLA loci.

## 2 Results

We use a publicly available data set of HIV-1 SPVL and HLA class I genotypes (HLA-A, B and C) from 327 patients from southern Africa [hiv.lanl.gov; 15]. After removing subjects for which the virus load is missing, or no HLA typing is available, we end up with 269 female patients of child-bearing age. The median log10 SPVL in this population is 4.47 log_10_ copies per ml blood (95% range: 2.60 – 6.06). From 87 patients, the complete HLA typing is known on the subtype level (2-field typing). The remainder of the patients have one or more loci typed at a type (1-field) or serotype level, or have one or more untyped loci.

We test a number of models, in order of increasing complexity, that predict SPVL, given HLA genotype (the models are explained in detail in the Methods section). In all models, we write a patient’s SPVL as a linear combination of weights *β* associated with her set of expressed HLA molecules:

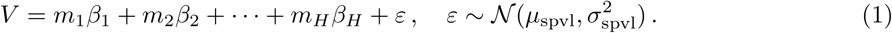

Here, the integer *m*_*h*_ϵ {0, 1, 2} indicates the copy number of the HLA allele *h*: *m*_*h*_ = 0 indicates that the patient does not have a particular HLA, while *m*_*h*_ = 1 and *m*_*h*_ = 2 indicate hetero and homozygosity for that HLA, respectively. The random variable *ε* represents the part of the SPVL that is not explained by HLA expression. The parameter *μ*_*spv1*_ is the average SPVL in this cohort and 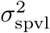 is the SPVL variance that is not explained by HLA.

Two models serve as control. First, a model that ignores HLA of the patient entirely, and models SPVL simply with a normal distribution, i.e. *V* ~ *ε* (the *null* model). The second model assumes independence of HLA alleles (the *independent* model). For each allele a single weight *β* _*h*_ is estimated, that are *a priori* assumed to be identical.

The *independent* model is similar to the models used in previous studies [19, 5], but differs in the sense that it is Bayesian. In previous (frequentist) approaches, HLA alleles that only occur in a small number of patients need to be excluded from the data, thereby reducing the number of parameters. Here we include all HLA alleles, but to keep the effective number of parameters in bounds, we assume that the weights *β* _*h*_ of the HLA alleles have a common prior distribution. We considered two different common prior distributions. First, the normal distribution with mean 0 and a standard deviation *σ* _branch_ is used. Note that *σ* _branch_ also needs to be estimated. This approach favors many small HLA effects, and relates to ridge regression in the frequentist framework, which is a penalized maximum likelihood method. Secondly, we also consider a Laplace prior distribution for the weights, which favors few, but possibly large HLA effects, and corresponds to the LASSO. To compare different models, we use the Watanabe-Akaike information criterion [WAIC; 44, 9], that is a measure for out-of-sample predictive power.

Another benefit of the Bayesian framework is that it provides a natural method for handling missing data. Often, patient HLA data contains missing values, due to difficulties in HLA imputation from raw data. HLA can be specified at multiple resolutions: single-field (e.g. A*02), two-field (e.g. A*02:01; any higher resolution has no effect on the amino-acid sequence of the HLA, and therefore on the binding groove), or an allele may be completely missing. For patients with single-field resolution HLA alleles or with any missing alleles, we sample 2-field alleles from all alleles within the same type or locus, taking population frequencies into account. For example, for a patient that has type B*58, we sample from B*58:01 and B*58:02 (see Methods). Notice that the issue of imputing high-resolution alleles can also be resolved in a frequentist framework, for instance by including “fractional” individuals [2], using *a priori* estimated probabilities of high resolution HLA expression [20]. As expected from previous studies, the *independent* model performs significantly better than the *null* model (ΔWAIC = 23.6, see Table 1). This simply asserts that SPVL depends strongly on HLA genotype.

The next models incorporate functional similarities between HLA molecules in several ways. In the Bayesian framework, we can encode these similarities as *a priori* information about the weights of the HLA alleles. The simplest way to accomplish this, is to choose *N*^*H*^ (0, σ) as a prior distribution for the weights, where σ is a covariance matrix based on the HLA similarities (we refer to these models as *multivariate normal* models). Notice that the *independent* model with a Normal prior distribution for the branch weights is characterized by σ = *σ* ^2^ · *I* with *I* the identity matrix. To characterize HLA similarity, we primarily use the non-consecutive amino-acids at 34 positions in the MHC molecule that are known to be variable, and in close proximity to the peptide binding groove (pseudo sequences). The amino-acids at these positions are provided by the peptide-binding predictor NetMHCpan [10, 28]

**Table 1:**
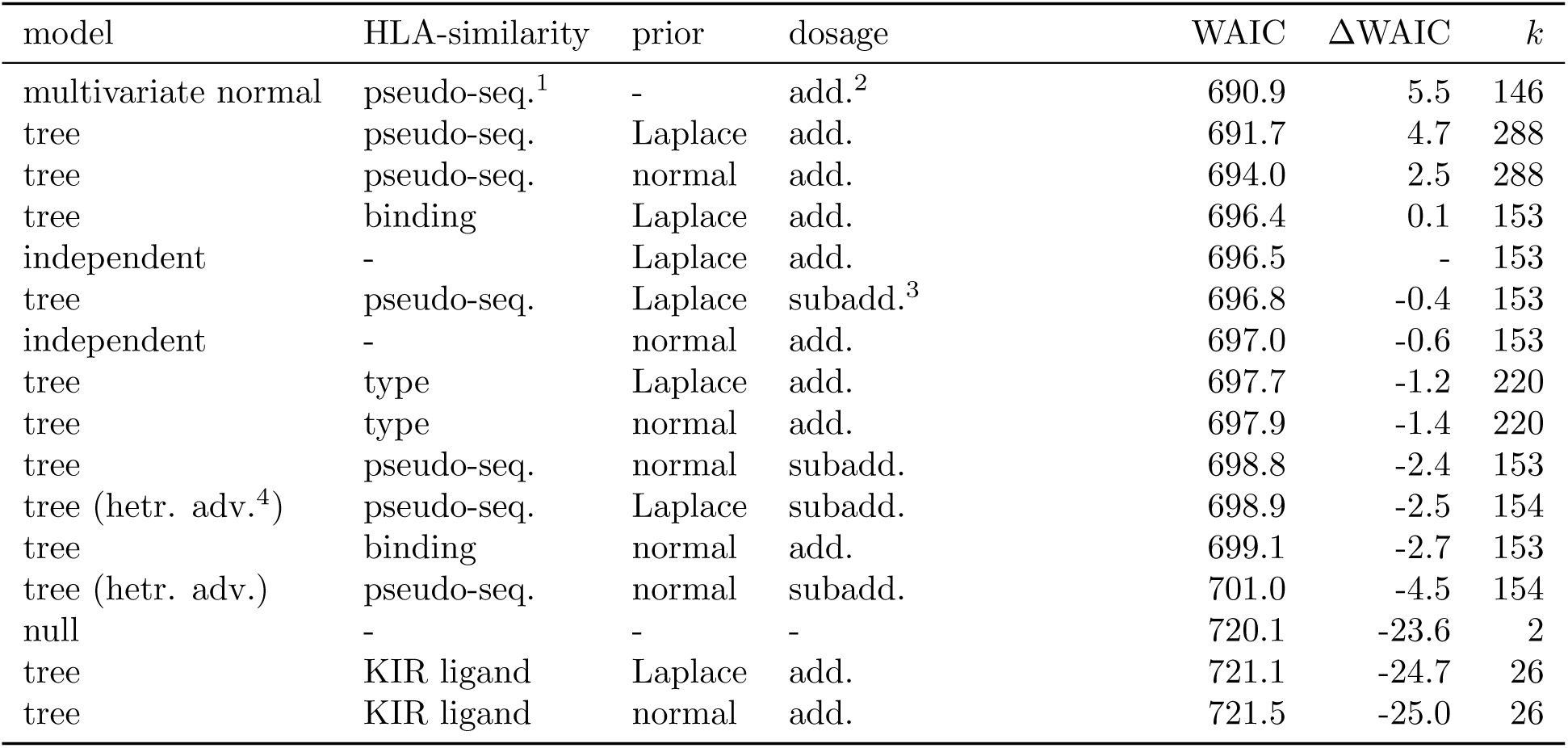
Comparing several models. The difference in WAIC (ΔWAIC) is taken with respect to the *independent* model. A lower WAIC indicates a better model fit, a ΔWAIC ≥ 2 is considered “positive” evidence for the better model, and a ΔWAIC ≥ 6 is considered “strong” evidence. The prior distribution for the edge weights is given in the *prior* column. The number of parameters (k) excludes those parameters used for estimating allele frequencies, missing alleles or censored SPVL. Notes: ^1^Pseudo-sequence. ^2^ Additive. ^3^ Subadditive. ^4^ Heterozygote advantage; a parameter (*η*) representing a generalized heterozygote advantage is added to the model.

**Figure 1:**
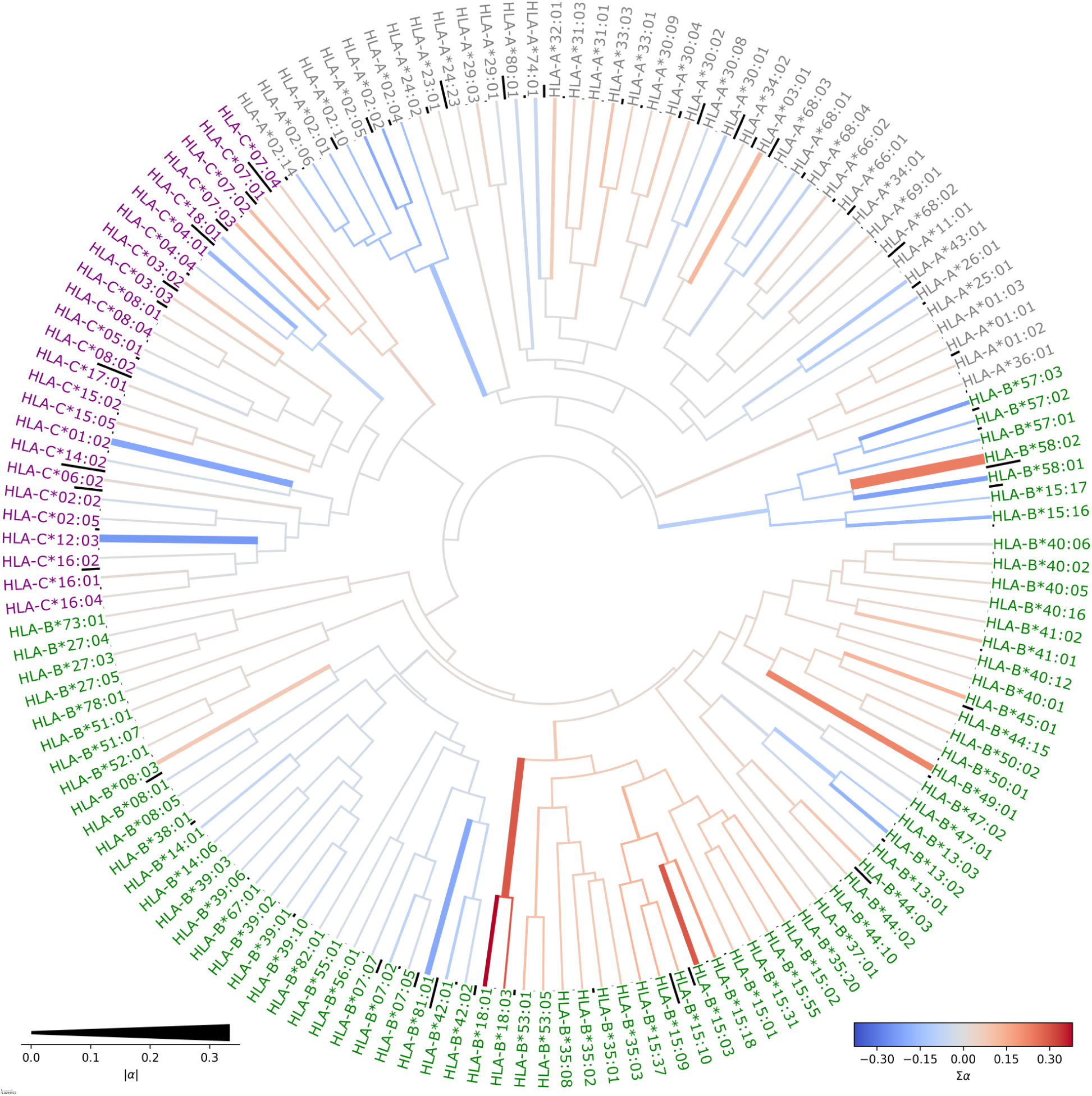
HLA tree with branches colored according to the estimated cluster-effect on SPVL. The HLA molecules are clustered using pseudo-sequences and the UPGMA algorithm. The color of the branches corresponds to the cumulative cluster effect (σ*α*) of the corresponding clade (protective: blue, detrimental: red). The thickness of the branches indicates the contribution (i.e. absolute weight, |*α*|) of the branch. The effect of thick lines is large (either protective or detrimental). The allele frequencies in the cohort are indicated by black bars at the leafs. Alleles of the loci A, B and C are colored gray, green and purple, respectively.

An alternative parameterization of the *multivariate normal* models is related to the phylogenetic mixed model [PMM; 11]. In the PMM, a quantitative trait *x* is assumed to have a genetic component *g*, and a random component *e* (hence, *x* = *g* + *e*). The genetic component *g* evolves according to Brownian motion on the branches of the phylogeny. SPVL can be considered a trait of the HLA molecules. As the phylogeny of the HLA molecules is rather complex, and does not encode functional diversity per se, we construct other hierarchical clusterings of the molecules using pseudo-sequences, peptide-binding predictions [43], or the nomenclature. Each branch of the clustering is given a weight *α*, and the weight of an HLA allele (a leaf in the tree) is defined as the sum of the weights *α* that is encountered by walking from root to leaf (see Figure 3). We refer to this model as the *tree* model. The weights *β* (Equation 1) are related to the weights *α* in accordance with Equation 3 in Methods.

If we cluster HLA molecules based on their pseudo-sequences, we find that the *multivariate normal* and *tree* model predict SPVL significantly better than the *independent* model (ΔWAIC = 5.5 and ΔWAIC = 4.7, respectively). surprisingly, the tree model is not favored over the independent model, when we cluster HLA molecules based on HIV-1 peptide-binding predictions (ΔWAIC = 0.1), indicating that the method is sensitive to the similarity measure.

As a predictive model, the *tree* model has no advantage over the *multivariate normal* model (ΔWAIC =-0.8), but it has some interesting benefits. The tree model allows us to color the HLA tree according to the weights of the branches (see Figure 1). This gives a quick overview into what level of resolution is responsible for any effect on SPVL. The most notable example is perhaps the B58 supertype, including the groups B*57 and B*58, and some alleles from the B*15 group. Most alleles in this supertype are protective, with the exception of the B*58:02 allele, which is associated with high SPVL. This property of B*58:02 is clearly visible in our *tree* model (Figure 1). The B*18 type, together with other alleles from the B*15 group appear to have a detrimental effect, while the A*02 group has an overall protective association. However, notice that the A*02 association could be largely driven by A*02:05, which is in linkage disequilibrium with the protective B*58:01 allele [15]. Notice that the model allows for estimation of supertype, type and subtype effects simultaneously.

The clusters in Figure 1 show a resemblance with the groups based on single-field typing—with some notable exceptions, e.g. B*15. These groups can be used to construct a simple HLA tree, that has one intermediate layer consisting of all single-field HLAs (HLA-A*01, HLA-B*07, etc.), and hence we can use our algorithm to estimate group-level weights, and subtype-level weights (Figure S2). The *type-tree* model made using this approach clearly uses some of the HLA type-level branches to capture the fact that molecules of the same type tend to be more similar. However, the predictive power of this type-based model is not superior to the *independent* model (ΔWAIC = −1.2; Table 1).

### 2.1 Predicting SPVL for out-of-sample HLA alleles

Until now, we have used the WAIC to compare the predictive performance of the models. Specifically, the WAIC tests how accurately a patients SPVL can be predicted, knowing the patients HLA genotype. The HLA molecules present in a single cohort are only a small subset of all existing, or even all known HLA molecules. When one considers two different populations, e.g. a European population and a Sub-Saharan African population, one will find roughly the same supertype distribution, but the HLA subtypes that represent the supertypes can be very different from one population to the next; e.g. for the A1 supertype, A*01:01 is most prevalent in Europe (28.6%), while A*26:01 is the most common representative subtype in Japan [38]. The estimated associations are therefore strongly dependent on the population that a cohort comes from [13]. Therefore, it is interesting to know how well our model predicts the contribution to SPVL of out-of-sample HLA alleles; i.e. HLA alleles that are not present in the cohort under investigation. For this, we use the leave-one-molecule-out approach [LOMO; 14],i.e. we train the model on all patients that do not express a selected HLA molecule, and then SPVL is predicted for the patients that do express that specific molecule.

As expected, the *tree* model outperforms the *independent* model for the majority of the HLA-A, B and C alleles, with respect to predicting virus load of patients expressing the out-of-sample allele (Figure 2; *P* = 0.009, *P* = 0.001, and *P* = 0.032 for HLA-A, B and C, respectively; Wilcoxon signed-rank test). An expected erroneous prediction is HLA-B*58:02, that is associated with fast progression, while the other alleles in the B*58 and B*57 groups are associated with low virus load. Therefore, training the *tree* model on all alleles except B*58:02, leads to the invalid prediction—based on the B*57/B*58 cluster effect—that B*58:02 must also be protective. The fact that the tree model gives erroneous predictions for the C*06:02 allele is due to linkage disequilibrium between B*58:02 and C*06:02 [15].

### 2.2 Generalizing the heterozygote advantage

Another benefit of the *tree* model is that we can easily change some of the assumptions. For now, we have ignored any effect of HLA homozygosity, which has been indicated as a cause of higher SPVL and faster disease progression [6]. In the models above, the contribution to the SPVL of two identical molecules is counted twice (see Methods). As an alternative scenario, we could count the weight at a homozygous locus only once. The HLA tree model allows us to generalize this idea. If identical molecules have a different effect than non-identical molecules, then perhaps also partially-identical molecules—as indicated by the functional HLA clustering—have a different effect on SPVL than two very distinct HLA molecules (see Figure 3). This idea is similar to the notion of gene dosage [45]. In Table 1, the *dosage* column is used to specify the method of counting weights of HLA molecules. *Subadditive dosage* indicates that the branch weights are counted only once.

The way the multiplicity of a branch is counted reveals another previously ignored problem concerning the biological interpretation of what it means to be a detrimental HLA allele. HLA molecules can be protective because they bind peptides that elicit a strong immune response, possibly inducing a costly escape mutation, or because of interaction with NK cells. The reasons why some HLA alleles are detrimental, however, are not as clearly understood. A simple explanation for finding alleles that are associated with high SPVL is that they are less protective than the average allele. More interestingly, some alleles appear to have a truly detrimental effect [27], possibly explained by the interaction between HLA-E expression and the NK-cell response [33]. The distinction between relative and true detrimental effects is important for the *subadditive dosage* model. In this model (see Methods, Equation 4) protective alleles correspond to negative branch weights, which are counted once if a patient is homozygous for an allele. Detrimental alleles correspond to positive weights, which means that patients who are homozygous for a detrimental allele are better off than heterozygous patients carrying two very different detrimental alleles. In other words, we here assume that detrimental molecules are truly detrimental in the sense that they are not merely less protective than the average molecule.

**Figure 2:**
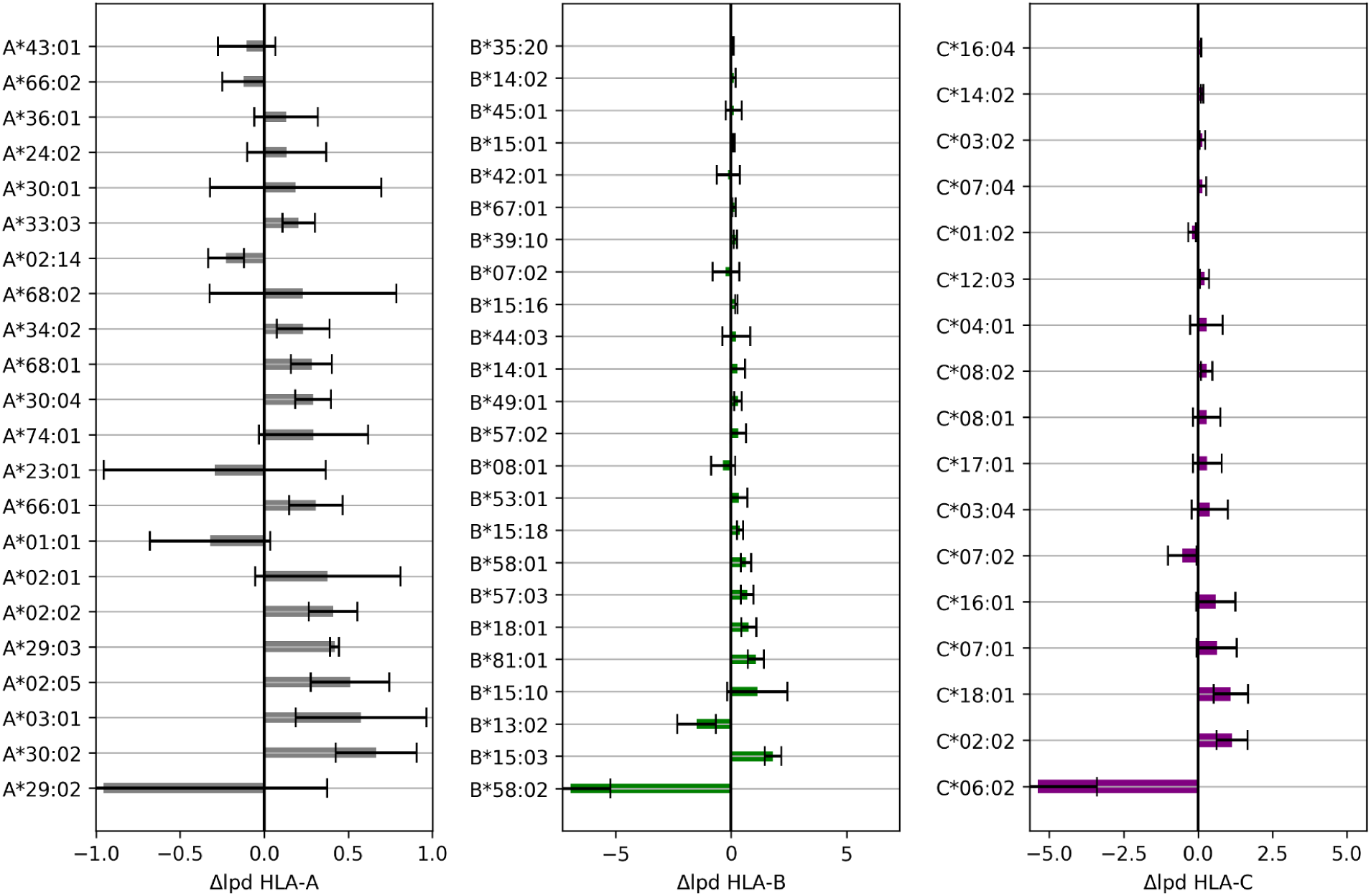
Leave-one-out cross-validation for the HLA alleles. For each HLA allele, a bar indicates the difference in log posterior density (Δlpd; see Methods). The error bars indicate the standard error. When Δlpd is positive, the *tree* model is better at predicting virus loads for patients with the out-of-sample HLA allele than the *independent* model. Only alleles with a Δlpd ≥ 0.1 are included in the figure.

For the *tree* model with *subadditive dosage* and a Laplace prior distribution on the branch weights, the resulting WAIC equals 696.8, indicating that the *subadditive dosage* model is significantly worse than the standard *tree* model (ΔWAIC = −5.1), and comparable to *independent* model. Therefore, we do not find evidence for a “generalized” heterozygote advantage. This is in agreement with findings of Leslie *et al.* [19], where it was suggested that the effect of HLA on SPVL is additive. However, as we have mentioned, other studies find evidence for a detrimental effect of HLA homozygosity on SPVL [6].

By adding a parameter to the linear model that can be interpreted as the generalized heterozygote advantage, we can model the situation in which all alleles are protective, and detrimental effects are simply relative (see Methods, Equation 6). However, based on WAIC, we can not distinguish between the model with truly (Equation 4) and relatively (Equation 6) detrimental HLA alleles (ΔWAIC = 0.1).

## 3 Discussion

We have trained Bayesian regression models, informed by the functional relatedness between HLA molecules, on publicly available HIV-1 SPVL and HLA genotype data. We found that a model that incorporates HLA similarities as prior information, outperforms a model that treats each allele as an independent predictor. Similar methods were presented by Sharon *et al.* [39] and Bartha *et al.* [1]. In particular, Bartha *et al.* [1] use the GCTA method [45], which is similar to our *multivariate normal* model. Bartha *et al.* [1] find that 8.4 ± 4% of the variance in SPVL can be explained by genetic variation in the MHC locus. Using a Bayesian equivalent of the coefficient of determination [8], we find a very similar value: 𝔼[*R*^*^2^*^] = 8%, 95% CrI: [3%, 14%].

Our method could still benefit from a number of improvements. First, the imputation of partially missing HLA allele data uses HLA allele frequencies and not HLA haplotype frequencies. It is known that linkage disequilibrium (LD) can be very strong between the B and C locus, and hence the accuracy of imputation could be improved by using haplotype frequency data. For more recent datasets with fewer missing data this would be less of an issue. LD between loci is also not explicitly incorporated in the estimation of the weights, although the linear model partially protects us from finding spurious associations. In fact, using the *tree* model could contain the LD problem even further. Suppose that two alleles are in LD, but their closely related neighbors on the tree are not. In this case an SPVL-association with the closely related allele can help to recover the true effect from the linked effect.

Secondly, the way we calculate the distances between HLA alleles can be improved by paying closer attention to the exact biological mechanisms that drive the associations. We have experimented with several distance measures, and found that a simple measure as the distance between pseudo-sequences derived from the HLA binding groove gives the best result. On the other hand, a more sophisticated method that uses the HIV-1 peptide-binding profiles of the HLA molecules does not outperform the *independent* model. This observation led us to hypothesize that perhaps other mechanisms than the CD8^+^ T-cell response are responsible for the observed associations.

To give an example: peptide-MHC complexes are not only ligands for the TCR, but also for the killer immunoglobilin-like receptor (KIR) of the natural killer (NK) cell. NK cells express a variety of receptors that are either activating or inhibiting. The ligands for inhibiting KIRs fall into five groups: HLA-A and B alleles with the Bw4 motif, the HLA-A3 and HLA-A11 serotypes, and the HLA-C1 and HLA-C2 groups. In order to see if a clustering of HLA alleles based on KIR epitopes can be used to explain HIV-1 SPVL, we constructed a HLA tree using the Bw4 motif, i.e., we use only five positions to define the functional distance between HLA molecules (see Methods and Figure S4).

The HLA tree resulting from the KIR-based distance metric contains only 12 leafs, which are representatives of 12 classes of HLA molecules. An identical effect is assumed for molecules in the same class. The estimated branch weights are very small compared to the weights estimated for the pseudo-sequence tree model (Figure 1). In fact, the KIR tree model is worse than even the null model, with a ΔWAIC of −1.1 (Table 1). Obviously, our KIR-inspired distance measure is very crude, since it does not take an individuals KIR genotype into account, let alone the KIRs that are expressed by a patients educated NK cells. Moreover, KIR-pMHC binding is not only determined by the Bw4 epitope, but also positions 7 and 8 of the presented peptide [42]. The fact that KIR genes are very polymorphic themselves suggests that a similar tree-based approach could be useful on the KIR-side of the analysis [23]. Not surprisingly, our KIR-based *tree* model can not explain any variation in the SPVL, it only adds complexity to the *null model*, without performing better.

For a third improvement to our model, we have to consider that the mechanisms that drive associations could be different for distinct clusters in the HLA tree. For instance, the association between B*57 and SPVL of HIV-1 could be due to CTLs and NK cells (or both), while the B*18 association could be caused only by CTLs. This would require different distance measures in different parts of the tree. Such an approach clearly needs further research to be implemented.

Research into the relationship between HIV-1 and HLA has moved far beyond identifying HLA associations with SPVL. In recent years, HLA has been identified as a driver of viral evolution [4], and even tumor evolution [24]. Viral polymorphisms can be linked to particular HLA alleles, types, and super-types. The importance of relatedness between HLA molecules has been appreciated in some of these studies [5, 2, 31]. For instance, Carlson *et al.* [2] found that differential CTL escape patterns within HLA groups and supertypes tend to be more common in more genetically diverse HLA groups. In a related study, Palmer *et al.* [31] systematically identify HLA-induced selection profiles on the HIV-1 genome, and use these to cluster HLA molecules. The resulting dendrogram is then compared to a HLA-protein derived dendrogram. They conclude that closely related HLA molecules have closely related selection profiles.

Because HLA associations with HIV-1 disease traits are so well established, we chose HIV-1 SPVL data to validate our method. For many diseases, HLA associations are not as clear as in the HIV-1 case, although we expect HLA to play a major discriminating role in many more diseases and in vaccine effectiveness. As the coeliac disease example shows [40], finding associations can be the first step in understanding the mechanisms behind a particular disease. Recently, Ramsuran *et al.* [33] discovered a mechanism that explains negative associations between HLA-A alleles and HIV-1 SPVL, and proposed a possible treatment exploiting this mechanism. Discovering HLA associations can thus ultimately lead to treatments and possibly cures. We believe that our method will help with discovering and understanding new HLA-disease associations.

## 4 Methods

In this paper, we consider a number of models. In each of these models, the log_10_ SPVL (indicated as *V* below) is determined partially by the presence of specific HLA class I molecules

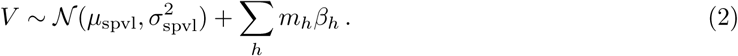

Here, *m*_*h*_ ϵ {0, 1, 2} is the multiplicity (or copy number) of HLA allele *h*, indicating absence (*m*_*h*_ = 0), and presence (*m*_*h*_ = 1 for heterozygous and *m*_*h*_ = 2 for homozygous). For each patient,Σ_*h*_ *m*_*h*_ = 6.

Equation 2 gives the likelihood of observing *V*, given weights *β* _*h*_, multiplicities *m*_*h*_, and the parameters *μ* _*spv1*_ and *σ* _*spv1*_ modeling random effects. Any information about relatedness of the HLA molecules is implemented in the model by specifying a prior distribution for the weights *β* _*h*_. In the case of the *tree* model, this prior is constructed using a relatedness-tree (*N, B*) with nodes *n* ϵ *N* and branches *b* ϵ *B* (Figure 3). The branches *b* are equipped with a length *δ*_*b*_ ≥ 0. Special nodes are the root *n*0 and the leafs *l*_*h*_, representing the HLA alleles. Each branch *b* is given a weight *α*_*b*_ with prior distribution 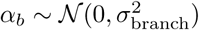 or *α*_*b*_ ˜ Laplace 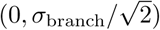 For each leaf *l*, a unique path (*n*_*0*_, *n*_*1*_, *n*_*2*_, *…, n*_*k*_ = *l*) exists that connects the root with the leaf. To determine the weight *β* _*h*_ of HLA allele *h*, we simply sum the weights *α* along this path, taking the branch lengths *δ* into account:

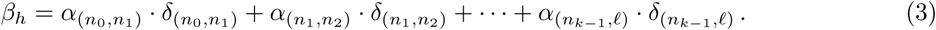

The weights *α* _*b*_ are estimated using Markov chain Monte Carlo (MCMC), using an Inv-Gamma(10^-3^, 10^-3^) hyperprior for the scale parameter 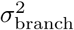. This hyperprior is also used for 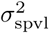, the standard deviation of the residuals of the log_10_ SPVL. The branch lengths *δ* _*b*_ are calculated using a clustering algorithm (see below).

### 4.1 Clustering HLA molecules

In order to construct the various HLA trees, we use the UPGMA algorithm with a HLA distance matrix as input. We use a number of different distances between two HLA molecules, depending on the similarity measure of interest. The branch lengths of all trees are scaled so that the height of the tree equals 1. The HLA trees can also be used to compute covariance matrices (Σ) that are used in the *multivariate normal* model. The covariance of two HLA molecules is equal to the depth (i.e. the distance to the root) of the “most recent common ancestor” of the molecules

**Figure 3:**
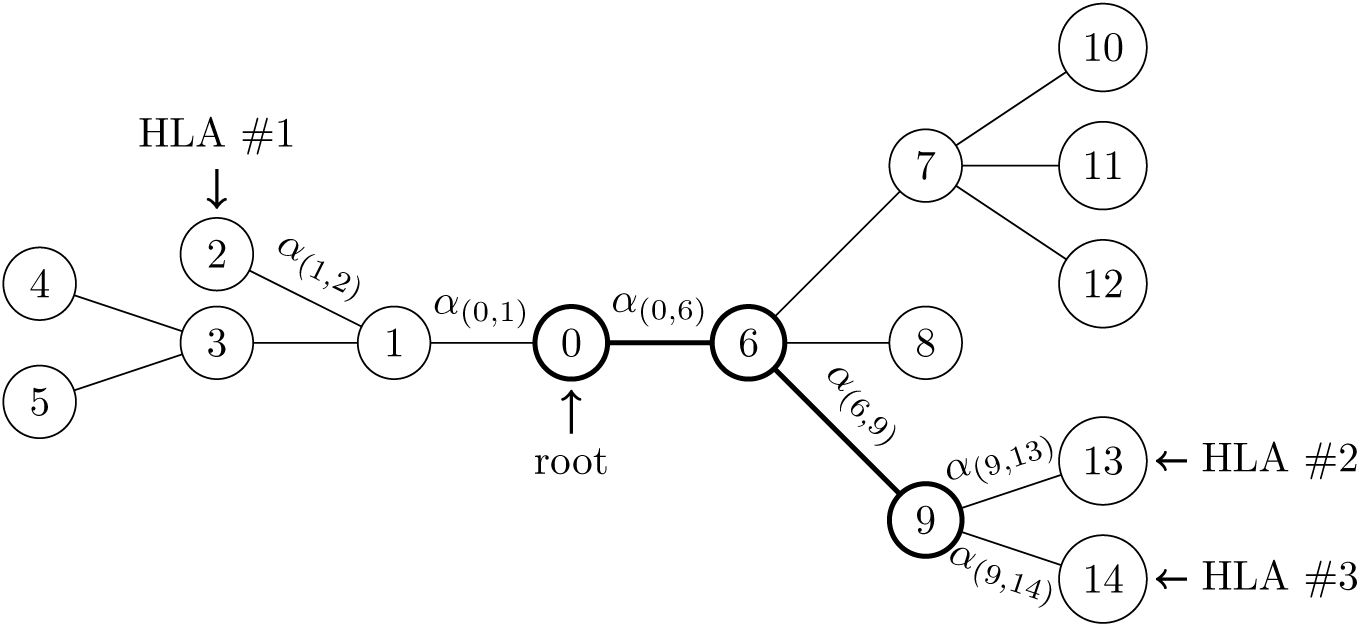
Example of a weighted HLA tree. The HLA molecule #1 corresponds to node 2, and the contribution to the SPVL of HLA #1 is given by *α*_(0,1)_ + *α* _(1,2)_Likewise, HLA #2 corresponds to node 13 and its contribution to the SPVL is *α* _(0,6)_+ *α* _(6,9)_+ *α* _(9,13)_For simplicity, in this example all branch lengths *δ* are equal to 1. In the *generalized homozygosity* model, the weights of the thick branches (0, 6) and (6, 9) are counted only once for a patient that carries both HLA #2 and HLA #3. In the default models, these weights are counted twice.

For the pseudo-sequence model, we use *n* = 34 amino-acids at positions 7, 9, 24, 45, 59, 62, 63, 66, 67,69, 70, 73, 74, 76, 77, 80, 81, 84, 95, 97, 99, 114, 116, 118, 143, 147, 150, 152, 156, 158, 159, 163, 167,171 of the HLA protein [exon 2 and 3, counting from the GSHSM motif; 29] We then use the PMBEC amino-acid covariance matrix *C* [16] to compute distances *d*_*i*_ = *d*(*a*_*i*_, *b*_*i*_) between two amino-acids *a*_*i*_ and *b*_*i*_ at position *i*. The covariance matrix is first converted into a centered matrix 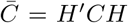 with *H*_*ab*_ = *δ* _*ab*_-1*/k*, with *δ* denoting the Kronecker delta. Here, *k* = 20 is the number of amino-acids. Then we transform the covariance matrix 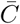 into a distance matrix using *d(a,b)*^2^ = (*k* − 1) 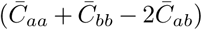. The distance between two HLA molecules is then defined as 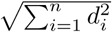

For the HLA tree that represents HLA similarities with respect to their role as KIR ligands, we use reduced psuedo-sequences (or motifs) of 5 amino-acids at positions 77, 80, 81, 82 and 83 of the HLA protein [22]. HLA protein sequences were downloaded from the IPD-IMGT/HLA database [www.ebi.ac.uk/ipd/imgt/hla; 36, 35].

Using this clustering approach can result in different molecules being indistinguishable from each other when they have identical pseudo-sequences. In those cases we only include a single representative leaf in the HLA tree, or a column in the covariance matrix, which needs to be non-singular for correct parameterization of the model. As we use only five positions to study KIR ligand effects on SPVL, many molecules were not distinguishable from each other, resulting in a tree with only 12 leafs. Allele frequencies are always estimated on the allele level, not merely on the level of the representative in the tree or covariance matrix.

For peptide-binding based clustering, we use the *MHCcluster* method [43], with the exception that only HIV-1 peptides are used to assess functional similarity. We downloaded a HIV-1 clade C reference sequence from South Africa from www.hiv.lanl.gov [18], and computed the binding affinity score for each 9-mer peptide from the proteome, using the binding-affinity predictor NetMHCpan (version 3.0). In addition to the binding affinity, NetMHCpan outputs a binding rank, that is calculated by comparing the binding affinity of a given peptide with the affinities of a large set of naturally occurring peptides. HLA molecules were then compared using Spearman’s correlation coefficient (*ρ*) between the binding-rank vectors. The correlations were converted into distances (*d*) using the relation *d*^*^2^*^ = 2 · (*n* −1) · (1 −*ρ*), where *n* is the number of peptides in the proteome.

### 4.2 Missing data

Some of the patients’ HLA typing is incomplete. Either a locus is missing, or an allele is only specified at the HLA-type level (1-field HLA typing). As we model HLA-similarity using pseudo-sequences or peptide-MHC binding predictions, the model requires 2-field HLA subtypes, and we model the missing HLA data as categorical parameters. Suppose that for individual *i* on locus *l ϵ* {A, B, C}, and gamete *u ϵ* {1, 2} the alleles *h*_*1*_, *h*_*2*_, *…, h*_*m*_ are admissible (i.e. the subtype belongs to the specified HLA type in the case of partial censoring, or the locus, in the case of complete censoring), then a priori *h*_*i,l,u*_ is drawn uniformly from *h*_*1*_, *h*_*2*_, *…, h*_*m*_.

In a population, some alleles are more common than others, and this fact should be used in the sampling of missing data [20]. We therefore model the HLA distribution *p*_*l*_ for the loci *l ϵ* {A, B, C}, and inform this distribution with the data form the cohort. Hence, given that we have sampled the missing HLA data from the admissible alleles, we multiply the posterior with ∏ _*i,l,u*_ *p*_*l*_ (*h*_*i,l,u*_). The allele frequencies *p*_*l*_ are estimated with Markov chain Monte Carlo (MCMC), together with the weights *α*. As a prior distribution we use *p*_*l*_ ~ Dirichlet(1*/*2, *…,* 1*/*2).

In addition to using the HLA frequencies in the cohort, we also use independent HLA frequency data from Sub-Saharan Africa (SSA) to further inform the HLA distribution [26]. This data consists of a vector *m*_*l*_ that sums to *M*_*l*_, and given *p*_*l*_, the observed frequencies *m*_*l*_ are Multinomial(*M*_*l*_, *p*_*l*_) distributed. Only the most common alleles, together covering at least 99% of the SSA dataset, were included in the analyses.

### 4.3 Leave-one-out cross-validation for HLA molecules

In order to test if the *tree* model can predict VL for out-of-sample HLA molecule better than the *independent* model, we perform for each HLA molecule *h* the following analysis. Patients that express the focal molecule *h* are excluded from the training set. However, in order to accurately estimate HLA frequencies, which is important for the sampling of missing HLA molecules, we still incorporate patients expressing *h* in the training set, but their SPVL is considered missing.

After fitting the *tree* and *independent* model to the partially masked data, we use both models to predict SPVL, and compare this to the observed SPVL. More precisely, we compute for each out-of-sample patient the log posterior density (lpd) of the observed SPVL, given the predicted SPVL and standard deviation *σ* _*spv1*_In Figure 2, we report the Δlpd; the difference between the lpd of the *tree* and the *independent* model, summed over all patients.

When a patient has (partially) missing HLA molecules, the left-out molecule *h* could be one of her admissible molecules, and in this case, we have to mask the SPVL of the patient. In the computation of the lpd for this patient, we then condition on her expressing the left-out molecules *h*, and the lpd is weighted by the posterior probability that the patient indeed expresses the molecule *h*. These posterior probabilities are also used to compute a weighted standard error of the Δlpd.

### 4.4 Subadditive dosage

In the standard *tree* model, each individual has 6 HLA molecules, corresponding to 6 paths in the HLA tree. The effect of the HLA molecules is given by the weights of the branches of each path. Each branch could (theoretically) occur 6 times in the linear model. This model assumes a dosage effect of HLA molecules; when a patient that is homozygous for a protective allele, the protective effect is assumed to count twice.

In the *subadditive dosage* model, the dosage effect is removed by counting each branch at most once. Hence, we have

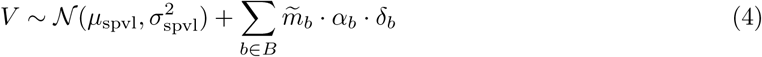

Where 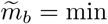{1, *m*_*b*_} and *m*_*b*_ ϵ {0, 1, *…,* 6} is the number of HLA molecules for which the branch *b* belongs to the corresponding path. The sum is taken over the branches of the tree. In this model, it is important how protective (negative weights) and detrimental (positive weights) branches are interpreted biologically.

Alternatively, one can assume that every molecule is protective. A hypothetical patient that lacks expression of HLA-A, B, and C molecules would have on average log_10_ SPVL equal to *V*_*max*_ [cf. 7]. Every expressed molecule *h* lowers the log_10_ virus load by 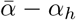, where 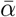is the average protective effect of an HLA allele, and *α* _*h*_ is the deviation from the expected value. This leads to the following model,

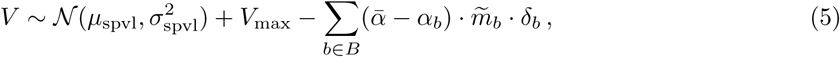

which is over-specified, and can be reduced to the following:

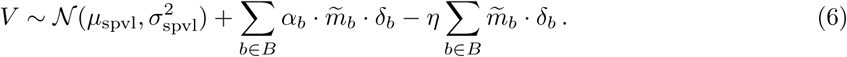

Here the quantity 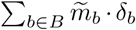 represents the level of heterozygosity of the patient, and the parameter *η* can be interpreted as the heterozygote advantage. The heterozygosity in the cohort equals 5.03 on average (2.5%-97.5% range [3.79, 5.47]).

**Figure S1:**
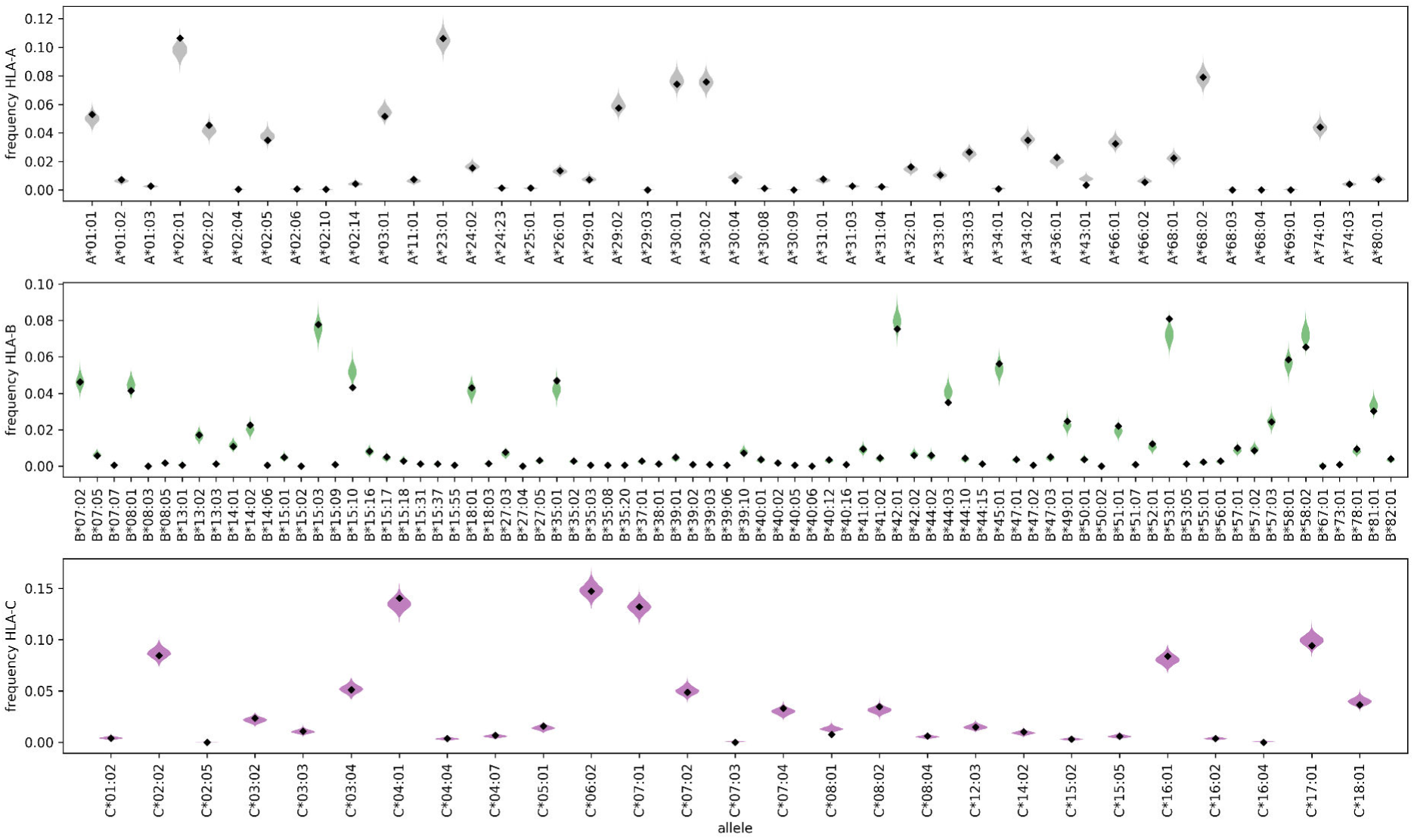
estimated HLA frequencies. The violin plots illustrate the posterior distribution of the HLA frequencies, using the *pseudo-sequence-tree* model with a Laplace prior for the branch weights. The diamonds give the point estimates based on the SSA population.

### 4.5 Implementation

The Bayesian models are implemented using JAGS [32], and the results are analyzed with Python. The trees are rendered with the ETE Toolkit [12]. A python module can be downloaded from www.github.com/chvandorp/MHCshrubs. For each model, we used chains of length 2 · 10^5^, discarding the first half as burn-in. Additionally, a 1: 100 thinning was applied to remove auto-correlation. Convergence of the chains was assessed by visual inspection of the trace plots of 4 parallel runs, and calculating the Gelman-Rubin statistic(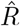 figureS5)

## 5 Acknowledgement

We gratefully acknowledge Jan van de Kassteele and Michiel van Boven for technical support with JAGS. This work is funded by The Netherlands Organisation for Scientific Research (www.nwo.nl, grant number 823.02.014).

---

**Figure S2:**
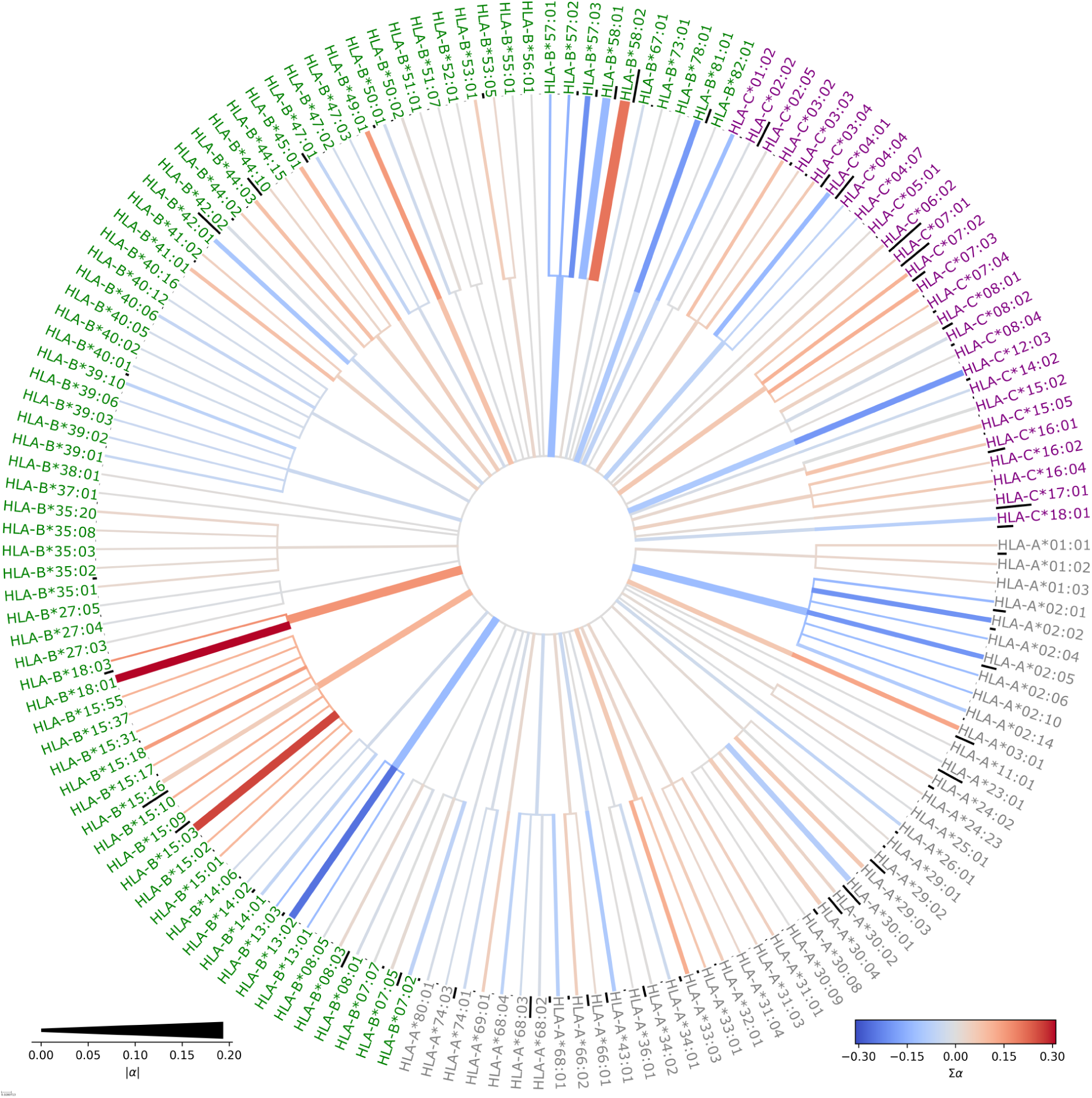
HLA tree based on single-field typing. Each HLA subtype (2-field typing) is represented by a leaf of the tree. Each HLA type (1-field typing) is represented by an intermediate node. The branch weights are given a Laplace prior distribution.

**Figure S3:**
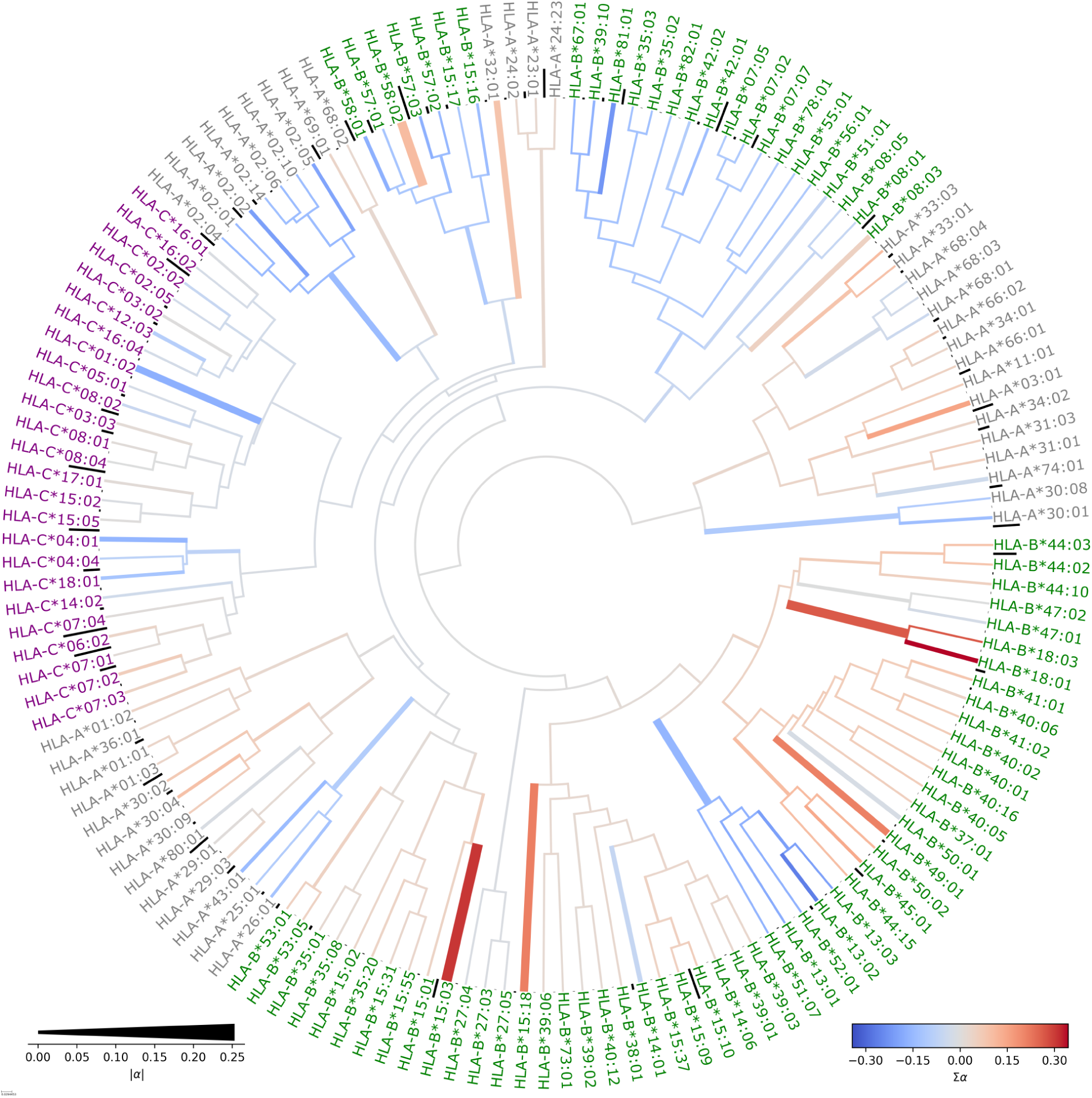
HLA tree based on peptide-binding predictions. For all 9-mers of a representative HIV-1 clade C proteome a binding rank is computed using NetMHCpan. The Spearman correlation between the binding ranks is used as a similarity measure between HLA molecules. The branch weights are given a Laplace prior distribution.

**Figure S4:**
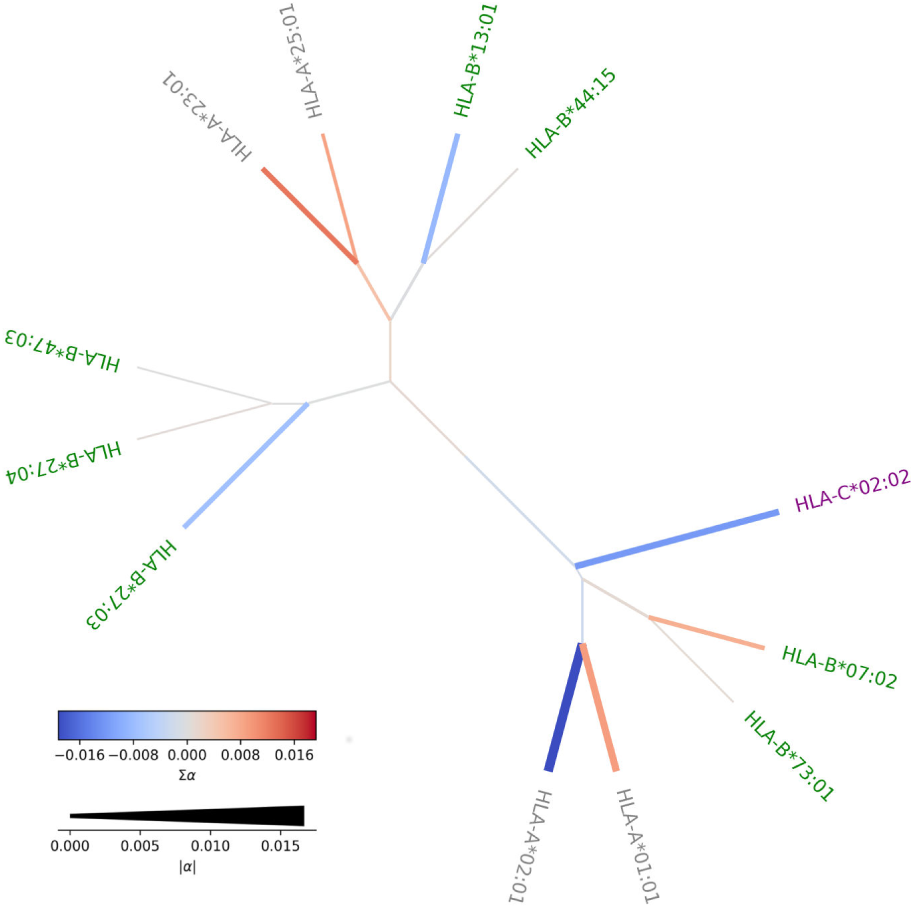
HLA-molecule clustering based on similarity as KIR ligands. The colors and widths of the branches indicate the contribution to the HIV-1 virus load, as in Figure 1, but notice the different scale.

**Figure S5:**
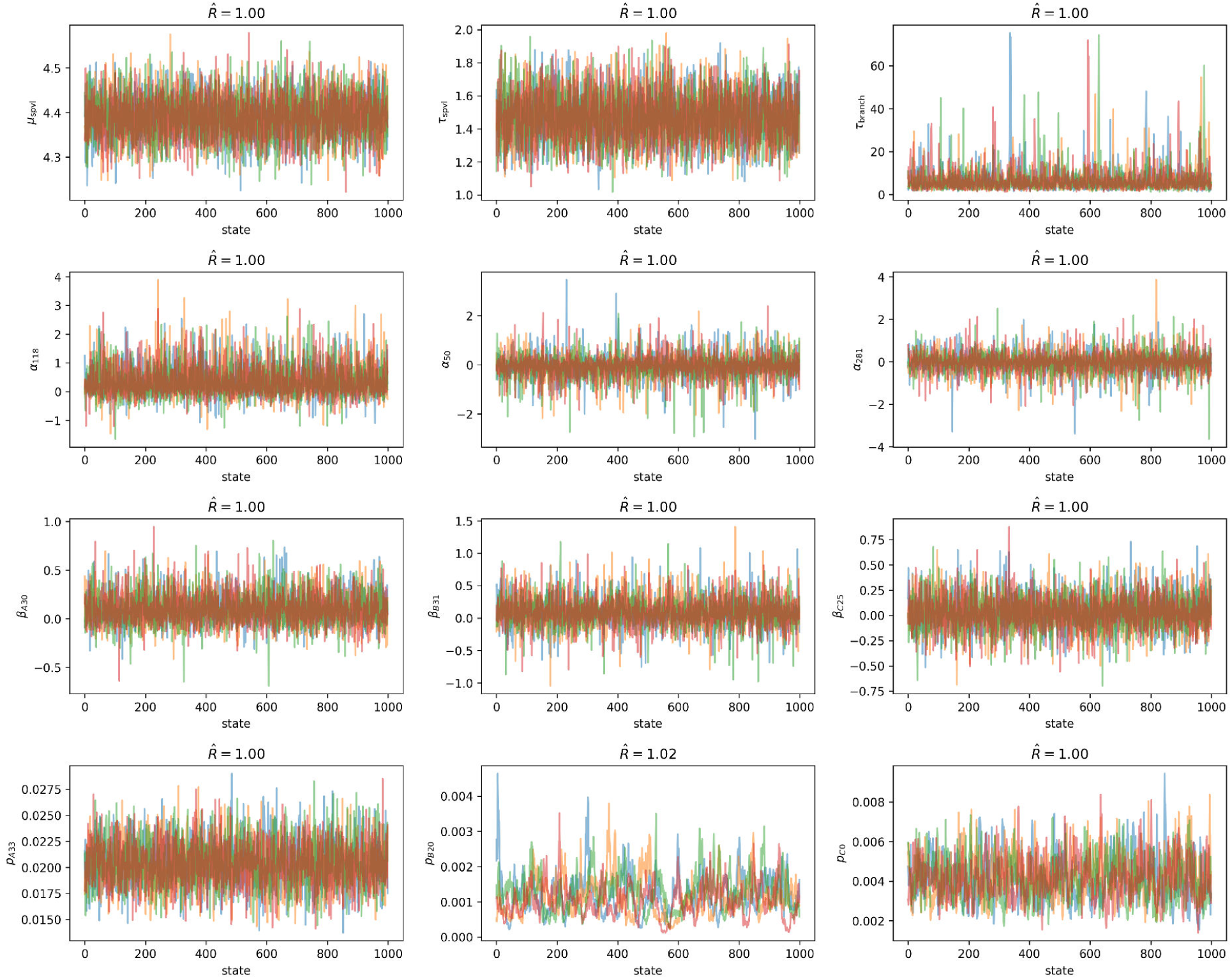
Examples of the traces of model parameters. Each panel shows the trace from 4 independent MCMC runs, and the Gelman-Rubin statistic 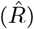. An 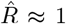 indicates convergence. Instead of the variance, the traces of the precision parameters are shown: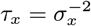 for *x* = spvl, branch.

